# Antimicrobial Resistance in Wildlife across Switzerland and the Principality of Liechtenstein: Associations with Environmental Factors and Taxonomic Variation

**DOI:** 10.1101/2025.01.28.635243

**Authors:** Katharina Höcketstaller, Iris A. Marti, Claudia Bank, Bahtiyar Yilmaz, Jens Becker

## Abstract

Antimicrobial resistance (AMR) presents a significant challenge to global public health. Addressing this challenge requires a comprehensive One Health approach that integrates efforts in human, veterinary, and environmental domains. Whereas AMR research in medical and veterinary domains is extensive, AMR research in wildlife has received less attention. However, AMR prevalence in the environment, and specifically in wildlife, ultimately affects human life. In a large study conducted in Switzerland and the Principality of Liechtenstein, we collected 410 rectal swab samples from 37 different wildlife species to quantify the prevalence of *Escherichia coli (E. coli)* and its resistance to 16 antimicrobial drugs, determined through phenotypical antimicrobial susceptibility testing.

The study yielded an 81.5% *E. coli* isolation rate with a considerable 10.8% of isolates resistant to one or more antimicrobial drugs. Notably, 4.5% of the isolates demonstrated multidrug resistance (MDR). Furthermore, we found that *E. coli* in omnivores exhibited the highest levels of AMR, significantly higher than in carnivores and herbivores. In addition to these dietary associations, we found that the percentage of forested areas surrounding the sampling locations was inversely related to AMR rates, suggesting that environmental conditions play a role in mitigating AMR.

This study provides a large-scale quantification of AMR in a broad range of wildlife species and also identifies diverse patterns of AMR in *E. coli,* highlighting variations associated with both host species and environmental conditions. Our findings emphasize the role of wildlife as potential indicators of environmental contamination with antibiotics and as reservoirs of resistant bacteria.

## 1. Introduction

Due to its far-reaching consequences on the ability to treat bacterial diseases in humans, animals, and the environment, the presence of drug-resistant bacteria represents a major worldwide public health challenge [1,2]. Research on antimicrobial resistance (AMR) is therefore a large, decade-old research field, which has gained even more attention following the World Health Organization’s 2023 publication of guidelines emphasizing the urgent necessity to establish and oversee strategies for AMR management [1]. In order to address AMR, a holistic, multidisciplinary approach known as One Health, is essential [3]. Whereas AMR research and monitoring in medical and veterinary settings is substantial and may continue to grow, AMR research in wildlife is still in its infancy [4]. Various studies that found AMR bacteria in wildlife species have highlighted the potential risk of wildlife serving as reservoirs of clinically relevant antimicrobial-resistant Gram-negative enteric bacteria [5,6,7,8,9,10]. Multiple studies have proposed an association between human activities and the prevalence of AMR in wildlife. This hypothesis is underpinned by the observation that wildlife populations living in near proximity to human settlements exhibit a higher incidence of AMR bacteria than those living in more isolated regions. This trend may be attributed to the increased exposure of these animals to anthropogenic factors such as pharmaceutical waste, agricultural runoff containing antimicrobial drugs, and direct interaction with human populations, which potentially facilitate the transmission and propagation of AMR pathogens [11,12]. A small set of studies has hypothesized that AMR prevalence might be linked to species-specific traits, such as feeding habits [11,13,14]. However, a comprehensive overview of AMR prevalence among various wildlife species in association with species-specific traits and habitat characteristics is lacking, since most studies have focused on specific wildlife species in predefined habitats [15,16].

We here present a large survey of AMR prevalence in *Escherichia coli* (*E. coli*) in multiple wildlife species. *E. coli* serves as an indicator species for AMR prevalence in wildlife microbiota, and allows for quantifying associations with species-specific traits and habitat characteristics. We hypothesized that the patterns of AMR prevalence in *E. coli* would be associated with species-specific traits of the host species, individual biological factors, and environmental characteristics of the host species’ habitat. To investigate this hypothesis, we determined the prevalence of AMR in *E. coli* across a diverse array of wildlife species in Switzerland and the Principality of Liechtenstein. Hence, this study provides a map of the distribution of AMR in wildlife and quantifies the correlation of interspecies differences and habitat conditions with the prevalence of AMR. This information is crucial for developing targeted strategies to mitigate the impact of AMR on wildlife, public health, and ecosystems.

## 2. Materials and methods

### 2.1 Study area and sample collection

From October 2022 to June 2023, wildlife samples were collected across Switzerland and the Principality of Liechtenstein. Before sampling, pre-labelled sampling kits were sent out to cantonal hunting authorities, researchers, and veterinarians. Kits contained sterile medium swabs (Heinz Herenz, Hamburg, Germany), disposable gloves, a small waterproof bag, instructions for swab collection, a prepaid envelope and a detailed questionnaire. Through the questionnaire, we gathered essential data including the coordinates of the collection site, date of sample acquisition, as well as the age, sex, weight, and time elapsed since death of the wildlife species. Detailed protocols for sample collection were meticulously outlined for participants through an instructional video and a summarized manual, ensuring the consistency and accuracy of the sampling process. Following collection, samples were required to be shipped under non-refrigerated conditions to an accredited laboratory within a 24-hour window, facilitating prompt and efficient processing for analysis.

Samples were obtained through three different approaches. First, veterinarians, game wardens and hunters collected samples from freshly hunted or recently deceased wildlife. Samples were taken within 24 hours of death. For this, no animal experiment permit was required, as samples were obtained from dead animals. Animals had been killed during hunts or activities of game wardens, or in accidents. Second, songbird (passerine) samples from living wild birds were collected by veterinarians of the Clinic for Ruminants, Vetsuisse Faculty, University of Bern during bird monitoring and ringing activities approved by the Swiss Federal Office for the Environment (FOEN) (authorization number: P38). Here, birds were caught in mist nets and gently placed in individual bags containing a removable sterile lining for transportation to the ringing facility. After removal of the birds from the bags, feces excreted by the birds were collected with sterile swabs from the lining by a veterinarian (last author). Third, live Eurasian lynx and European wildcats captured and anaesthetized in the framework of ongoing research and translocation projects. They were sampled by veterinarians of the Institute for Fish and Wildlife Health, University of Bern (animal experimentation permit BE61/2020/32505).

### 2.2 Isolation and identification of *E. coli* strains

Rectal swabs were processed by streaking the swabs onto chromogenic coliform agar plates (Oxoid, Wesel, Germany) and incubating the plates at 35±2°C for 21±3h. For further characterization and species identification, dark blue or purple grown colonies suspected to be *E. coli* were specifically picked. Distinct colonies were carefully selected and transferred onto BD Columbia III 5% sheep blood agar plates (BD, Heidelberg, Germany) using the same incubation conditions and parameters. Species identification of the isolates was performed using matrix-assisted laser desorption/ionization time-of-flight mass spectrometry (MALDI-TOF, Bruker, Bremen, Germany). Once identified as *E. coli*, one bacterial isolate per sample was stored in Microbank vials at -20°C (Pro-Lab Diagnostics, Richmond Hill, Ontario, Canada). After shipping to the Vetsuisse Faculty, University of Bern, Microbank vials were stored at -80°C until further use.

### 2.3 Antimicrobial susceptibility testing

Antimicrobial susceptibility testing was performed using the disk diffusion method recommended by the Clinical and Laboratory Standards Institute (CLSI) [17]. Sixteen antimicrobial drugs from ten different classes were selected based on their relevance to human and veterinary medicine. Antimicrobial classes tested were (μg/disc): tetracyclines [tetracycline (30)], sulfonamides [trimethoprim-sulfamethoxazole (1.25/23.75)], penicillins [ampicillin (10), amoxicillin-clavulanic acid (20/10)], cephalosporins [cephazolin (30), ceftriaxone (30), cefotaxime (30), cefepime (30)], phenicols [chloramphenicol (30)], aminoglycosides [gentamycin (10), streptomycin (10)], fluoroquinolones [ciprofloxacin (5), nalidixic-acid (30)], glycyclines [tigecycline (15)], macrolides [azithromycin (15)] and carbapenems [meropenem (10)] (Marnes-la-Coquette, France). As control strain *E. coli* ATCC 25922 was used (Remel Inc., Lenexa, USA).

Each *E. coli* isolate was cultivated on brain-heart infusion agar plates (BHI) supplemented with 5% sheep blood (Bio Rad, Marnes-la-Coquette, France) and incubated at 35±2 °C overnight to facilitate growth. Subsequently, colonies were transferred to a sterile broth to achieve a 0.5 McFarland standard turbidity. The standardized bacterial suspension was evenly distributed onto Mueller-Hinton agar plates (MH) using a sterile cotton swab. Then, disks containing antimicrobial drugs were positioned on the inoculated MH agar plates using a disk dispenser (Bio Rad, Marnes-la-Coquette, France). After aerobic incubating for 16-18 hours at 35±2°C, the diameter of each zone of inhibition was measured using a calibrated ruler. Interpretive criteria by CLSI were used to classify isolates as susceptible (S), intermediate (I), or resistant (R) [18]. For Tigecycline, interpretive criteria of the European Committee on Antimicrobial susceptibility Testing (EUCAST) were applied since no CLSI breakpoints were available [19]. In this study, isolates categorized as intermediate were deemed susceptible, leading to the definition ‘AMR’ being used only to describe isolates that exhibited resistance. Multidrug resistance (MDR) was defined as the resistance of an isolate to at least one antimicrobial drug of three or more distinct classes of antimicrobials [20].

### 2.4. Evaluation of the geographical coordinates

Additional data concerning topography and habitat of the sample locations was provided by experts for governmental databases of the Federal Office of Topography (swisstopo, Wabern, Switzerland) and the Federal Office for Agriculture (FOAG). Data were retrieved for a one-kilometer radius around each sample location, and the following factors were recorded: altitude above sea level, population density in people per hectare (%), the proportion of forested area (%), the proportion of agricultural land (%), and the proportion of area of water bodies (%) as well as length of water bodies.

Altitude data was retrieved using ArcMap Tool function “extract values to points” using dataset SwissALTI3D 2m (swisstopo, Wabern, Switzerland). Population density was retrieved using ArcGIS Pro Tool “Zonal Statistics as Table” per 100mx100m plots (Federal Statistical Office, Neuchâtel, Switzerland). Percentage of forested area was retrieved using swiss TLM3D (swisstopo, Wabern, Switzerland) and Arc GIS Pro Tool “Tabulate Intersection”. Water bodies were assessed as water surfaces ≥5m width. Length of water bodies was assessed for all water bodies in a 1km radius for each sample location using the above-mentioned tools.

### 2.5 Statistical analysis

Data were handled using Microsoft® Excel (Microsoft, Redmond, USA) and statistical analyses were conducted using *lme4* package in R (R version 4.2.1, Vienna, Austria). The unit of the analysis was the bacterial isolate (one per wild animal). Associations between the susceptibility status of the bacteria as binary outcome (presence of resistance against tested drug(s) vs. pansusceptibility). Each potential predictor was first analyzed in a univariate logistic regression model. Potential predictors are shown in Table 1. Only potential predictors significant at P ≤ 0.05 that showed no collinearity were considered for further analyses. Potential predictors were considered as correlating when the mean square contingency coefficient ϕ was >0.6.

**Table 1.** Results of the logistic regression model for *Escherichia coli* isolate from wildlife in Switzerland and the Principality of Liechtenstein exhibiting resistance to ≥1 of the tested antimicrobial drugs.

| *Predictors* | *OR* | *CI* | *p* |
| --- | --- | --- | --- |
| (Intercept) | 0.08 | 0.02 – 0.22 | **<0.001** |
| Percentage forested area | 0.98 | 0.96 – 1.00 | **0.027** |
| Feeding behavior [carnivore, ref: herbivore]] | 1.08 | 0.15 – 5.8 | 0.929 |
| Feeding behavior [omnivore] | 4.6 | 1.74 – 15.92 | **0.006** |

To model presence and absence of pansusceptible isolates, binomial general linear models were fitted to the data using the following formula:

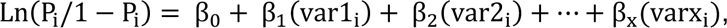

where P_i_ is the probability of being a pansusceptible isolate, β_0_ is the intercept from the linear regression equation, and β_1_ to β_x_ are the regression coefficients corresponding to the independent predictor variables var1_i_ to varx_i_ [21]. Odds ratios and confidence intervals were calculated from the model estimates. The model assumptions (linearity of independent predictors and log-odds, and absence of multicollinearity) were tested using Box-Tidwell test and variance inflation factor (VIF), respectively. The model’s significance level was set at p ≤ 0.05.

## 3. Results

### 3.1 *E. coli* in wildlife samples

A total of 410 rectal swab samples were collected from wildlife across Switzerland and the Principality of Liechtenstein (Figure 1). The sampled population consisted of 24 bird species (n=83) and 13 mammal species (n=251, Table 2 and 3). From these, 334 *E. coli* isolates were obtained (overall isolation rate: 81.5%).

**Fig. 1.**
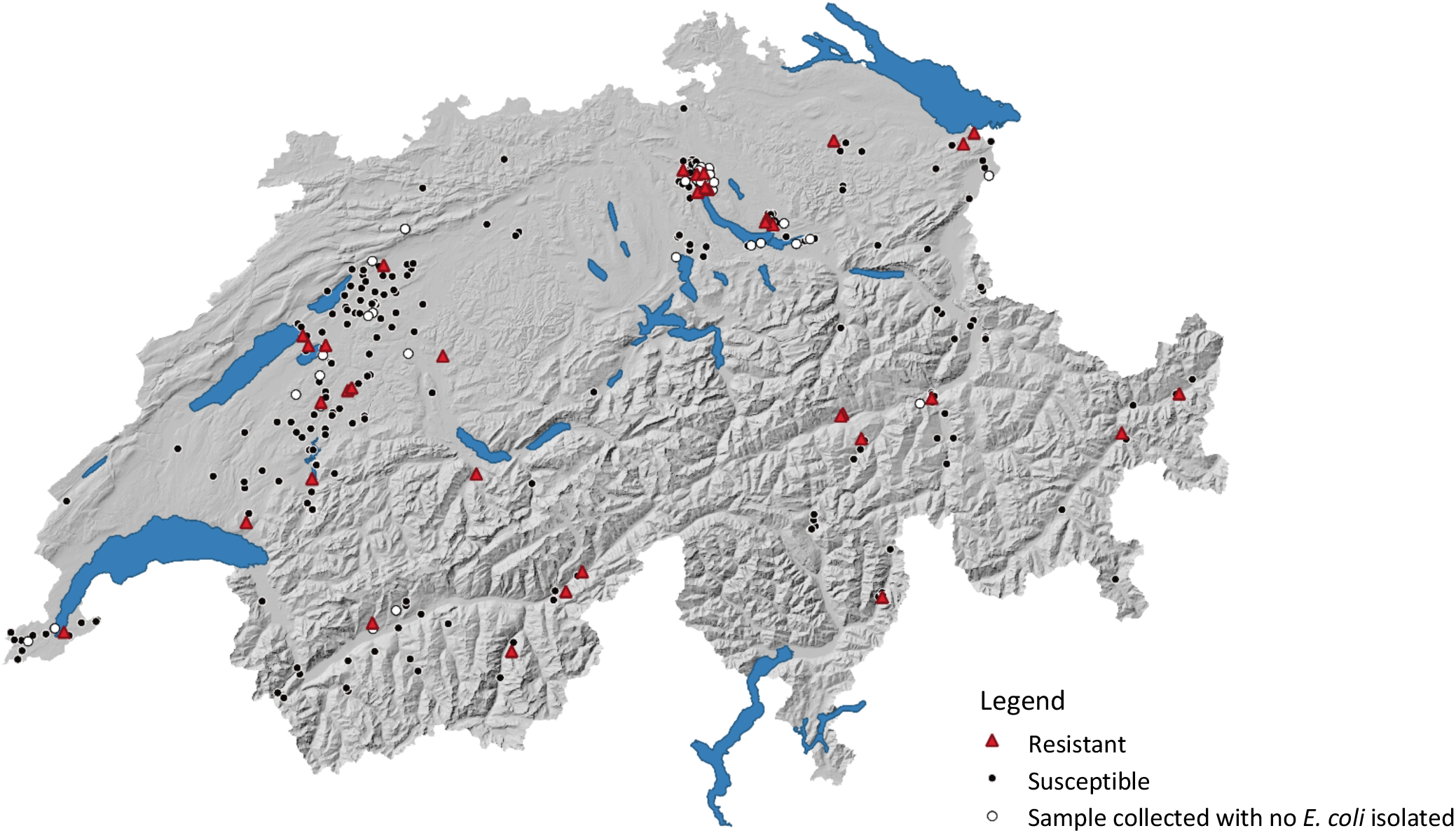
Locations of the collected samples of wildlife in Switzerland and the Principality of Liechtenstein collected from October 2022 to June 2023.

**Table 2.** Species of sampled mammals (n=251) and their feeding behavior, frequency of *Escherichia coli (E.coli)* isolation as well as number and prevalence of isolates exhibiting ≥1 resistance to the tested antimicrobial drugs in wildlife grouped by species.

| Common name<br>(*scientific name*) | Feeding<br>behavior | No. resistant<br>Isolates (%) | No. *E. coli*<br>isolates |
| --- | --- | --- | --- |
| Red fox (*Vulpes vulpes*) | Omnivore | 12 (14.6) | 82 |
| European badger (*Meles meles*) | Omnivore | 6 (27.3) | 22 |
| Wild boar (*Sus scrofa*) | Omnivore | 3 (12.0) | 25 |
| European roe deer (*Capreolus capreolus*) | Herbivore | 2 (2.53) | 79 |
| Eurasian lynx (*Lynx lynx*) | Carnivore | 1 (11.1) | 9 |
| Alpine chamois (*Rupicapra rupicapra*) | Herbivore | 1 (12.5) | 8 |
| European wildcat (*Felis silvestris*) | Carnivore | 0 (0.0) | 6 |
| Gray wolf (*Canis lupus*) | Carnivore | 0 (0.0) | 2 |
| European red deer (*Cervus elaphus*) | Herbivore | 0 (0.0) | 10 |
| European beaver (*Castor fiber*) | Herbivore | 0 (0.0) | 1 |
| Brown hare (*Lepus europaeus*) | Herbivore | 0 (0.0) | 1 |
| Red squirrel (*Sciurus vulgaris*) | Omnivore | 0 (0.0) | 3 |
| Stone marten (*Martes foina*) | Omnivore | 0 (0.0) | 3 |

The isolation rate varied significantly among the wildlife species (median: 81.5%; range: 37.3%-98.4%). The isolation rate was lowest in the bird population (57.6%), ranging from passeriformes (37.3%) to raptors (87.5%). The highest isolation rate was observed in artiodactyla (98.4%).

### 3.2 Antimicrobial susceptibility of *E. coli*

A total of 10.8% (n=36) isolates were found to be resistant to at least one antimicrobial drug. Isolates exhibiting AMR to ampicillin were most prevalent among the resistant isolates (66.7%; n=24), followed by cefazolin (55.6%; n=20), tetracycline (50.0%; n=18), and streptomycin (38.9%; n=14). Susceptibility to all tested antimicrobial drugs is shown in Figure 2.

**Fig. 2.**
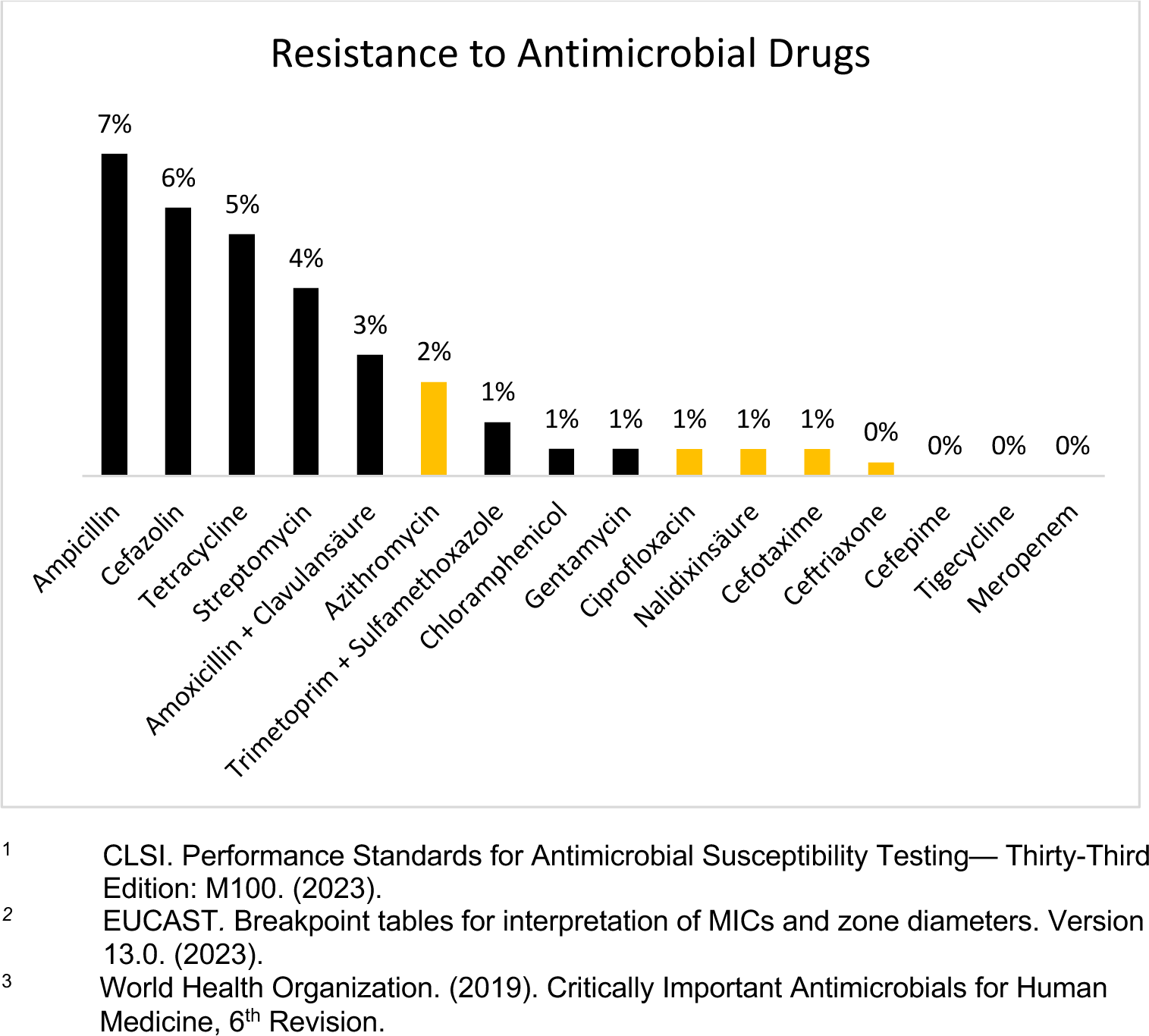
Antimicrobial drugs 334 *E. coli* isolates from wildlife exhibited resistance to. Interpretive criteria were retrieved from CLSI^1^ and EUCAST^2^. Samples were collected from birds and mammals across Switzerland and Liechtenstein. Antimicrobial drugs listed by the WHO as critically important antimicrobials^3^ are highlighted in yellow.

Notably, 3.0% of the isolates, corresponding to 10 samples, showed resistance to antimicrobial drugs that the World Health Organization categorizes as critically important antimicrobials (CIAs), as outlined in their 2019 report [22]. These CIAs encompass a range of vital antimicrobial classes including macrolides, fluoroquinolones, and third- and fourth-generation cephalosporins. Further analysis of multidrug resistance (MDR) revealed that 41.7% of the 36 AMR *E. coli* isolates (i.e., 15 isolates) displayed 12 distinct MDR phenotypes, as detailed in Table 4.

**Table 3.** Species of sampled birds (n=83) and their migratory behavior, frequency of *Escherichia coli (E. coli)* isolation as well as number and prevalence of isolates exhibiting ≥1 resistance to the tested drugs in wild animals grouped by species.

| Common name<br>(*scientific name*) | Migratory<br>behavior | No. resistant<br>Isolates (%) | No. *E. coli*<br>isolates |
| --- | --- | --- | --- |
| Mallard (*Anas platyrhynchos*) | Resident to<br>short distance | 3 (27.3) | 11 |
| Carrion crow (*Corvus corone*) | Resident to<br>short distance | 2 (22.2) | 9 |
| Black-headed gull (*Larus ridibundus*) | Resident to<br>short distance | 2 (50.0) | 4 |
| Feral pigeon (*Columba livia f. domestica*) | Resident | 1 (14.3) | 7 |
| Mute swan (*Cygnus olor*) | Resident | 1 (33.3) | 3 |
| Mandarin duck (*Aix galericulata*) | Resident | 1 (50.0) | 2 |
| Eurasian buzzard (*Buteo buteo*) | Resident to<br>short distance | 1 (20.0) | 5 |
| House martin (*Delichon urbicum*) | Long distance | 0 (0.0) | 1 |
| Common redstart (*Phoenicurus phoenicurus*) | Long distance | 0 (0.0) | 1 |
| Common raven (*Corvus corax*) | Resident | 0 (0.0) | 1 |
| Eagle owl (*Bubo bubo*) | Resident | 0 (0.0) | 1 |
| Egyptian goose (*Alopochen aegyptiaca*) | Resident | 0 (0.0) | 1 |
| Eurasian woodcock (*Scolopax rusticola*) | Resident | 0 (0.0) | 1 |
| Blackbird (*Turdus merula*) | Resident to<br>short distance | 0 (0.0) | 5 |
| Red kite (*Milvus milvus*) | Resident to<br>short distance | 0 (0.0) | 2 |
| Common coot (*Fulica atra*) | Resident to<br>short distance | 0 (0.0) | 1 |
| Rook (*Corvus frugilegus*) | Resident to<br>short distance | 0 (0.0) | 1 |
| Mediterranean yellow-legged gull (*Larus*<br>*michahellis*) | Resident to<br>short distance | 0 (0.0) | 1 |
| Great cormorant (*Phalacrocorax carbo*) | Short distance | 0 (0.0) | 17 |
| Blackcap (*Sylvia atricapilla*) | Short distance | 0 (0.0) | 3 |
| Chiffchaff (*Phylloscopus collybita*) | Short distance | 0 (0.0) | 2 |
| Grey heron (*Ardea cinerea*) | Short distance | 0 (0.0) | 2 |
| Black redstart (*Phoenicurus ochruros*) | Short distance | 0 (0.0) | 1 |
| European robin (*Erithacus rubecula*) | Short distance | 0 (0.0) | 1 |

**Table 4.** Phenotypic MDR profiles for antimicrobial drug combinations to which Escherichia coli (*E. coli)* isolates were resistant. Samples were collected from birds and mammals across Switzerland and Liechtenstein.

| Antimicrobial drugs | Species | No. *E. coli* isolates |
| --- | --- | --- |
| AMP-CZN-AMC-AZM | Wild boar, Red fox,<br>Mallard | 3 |
| AMP-CZN-TET-SMN | Black-headed gull, Wild<br>boar | 2 |
| AMP-CZN-TET | European badger | 1 |
| AMP-SMN-STX | Mute swan | 1 |
| AMP-TET-SMN-STX | Mallard | 1 |
| AMP-TET-SMN-AZM | Red fox | 1 |
| AMP-CZN-TET-AZM | Red fox | 1 |
| AMP-TET-SMN-AMC-STX | European badger | 1 |
| AMP-CZN-TET-SMN-CIP-NAL | Mallard | 1 |
| AMP-CZN-TET-SMN-CHL-GEN | Red fox | 1 |
| AMP-CZN-TET-SMN-CTX-CRO | Black-headed gull | 1 |
| AMP-CZN-TET-AMC-STX-CHL-CIP-CTX-NAL | Red fox | 1 |

AMC – amoxicillin/clavulanic acid; AMP – ampicillin; AZM – azithromycin; CHL - chloramphenicol; CIP – ciprofloxacin; CRO – ceftriaxone; CTX – cefotaxime; CZN – cefazolin; FEP – cefepime; GEN – gentamycin; MEM – meropenem; NAL – nalidixic acid; SMN – streptomycin; STX – sulfamethoxazole/trimethoprim; TET – tetracycline; TGC – tigecycline.

### 3.4 Association of resistant *E. coli* isolates with topographical and taxonomic data

Wildlife grouped by species, feeding behavior, and migratory behavior are shown in Table 2 and Table 3. For mammals, we grouped the wildlife species depending on feeding behavior in herbivorous species, carnivores and omnivores. For birds, we grouped the examined species due to their migratory behavior. Our study found a significant association between feeding behavior of mammals and AMR prevalence, as well as a significant association between higher overall AMR prevalence and lower levels of forested areas in the sample location, as shown in Table 1. Our analysis revealed no significant associations between AMR prevalence and individual factors such as age, weight, and sex among the sampled wildlife. Additionally, environmental factors, including the altitude of the sampling location, human population density, the extent of agricultural land, and the area covered by water bodies, did not exhibit a significant association with AMR prevalence in the isolated *E. coli*.

## 4. Discussion

This large-scale study, which examined a large number of different wildlife species from across Switzerland and Liechtenstein, is the first local study to comprehensively investigate the associations between wildlife, their species-specific traits, and habitat characteristics with the prevalence of AMR. Our findings reinforce the notion that the natural habitats of wildlife and their unique behaviors, particularly in terms of feeding, are key determinants in the prevalence of AMR. In addition to validating previous hypotheses and findings in AMR research, our study highlights the association of environmental characteristics with the patterns of AMR. The presence of MDR in such a significant proportion of resistant isolates stresses the concerning trend of increased antimicrobial resistance, particularly when considering the critical nature of the antimicrobials involved.

The observed level of AMR prevalence in *E. coli* isolated from Swiss wildlife was lower than previously reported AMR levels of wildlife from other European countries. However, some of these previous studies used different methods for susceptibility testing with different antimicrobial drugs, and their sampling methods varied. In Italy, observed AMR prevalence in various wildlife species ranged from 33.9% to 89.2% [16,23,24]. Similarly, studies conducted in Portugal, Canada, and England, also showed notably higher AMR prevalence, ranging from 23.0% to 60.8% [25,26,27]. However, other studies reported levels of AMR prevalence below 10.0% [6,8,13]. Notably, two of these studies were carried out in Germany, which is located in geographical proximity to Switzerland and the Principality of Liechtenstein is likely that the different observed levels of AMR prevalence in wildlife are connected to the country-specific AMR prevalence and antimicrobial drug consumption in humans and welfare. For example, the EFSA report showed significant differences in AMR prevalence in livestock in Europe, indicating a north-south gradient with higher values in the south [28]. In addition, the overall consumption of antimicrobial drugs tends to be higher in southern European countries than in northern countries [29,30]. Together, this suggests that higher AMR prevalence in wildlife aligns with increased antimicrobial consumption, reinforcing a call for a holistic monitoring approach and for including wildlife in considerations of potential AMR refugia and spillover events.

Since sampling methods, laboratory protocols used for susceptibility testing, distribution and number of sampling locations and sampled wildlife species show great variability among the cited international studies, caution is warranted when comparing study results. Here, the use of standardized test procedures could facilitate comparability, as well as careful consideration of habitats and different wildlife species. A single study from 2019 focused on Swiss wild birds that were admitted to a surveillance and rescue center with no information regarding the bird’s origin [31]. The described AMR prevalence and the sampled bird species are consistent with our findings and the bird species tested in this study.

In this study, we screened for resistance to the most frequently used antimicrobial classes in hospital care in human medicine in Switzerland[3], and the most frequently used antimicrobial classes in veterinary medicine in European countries [30]. Our study reveals an association between the prevalence of AMR in wildlife and commonly used antimicrobial classes mentioned above. This association aligns with findings from diverse studies conducted in other European countries [6,11,32], strengthening the suggestion of diverse transmission pathways of AMR bacteria from humans and domestic animals to wildlife [9]. This suggestion is further supported by two studies that investigated the influence of shared environments as a possible transmission pathway towards AMR prevalence in wildlife [24,33]. The association found in these two studies between shared environments and AMR prevalence among wildlife and livestock, as well as between wildlife, livestock, and humans, underlines that transmission to wildlife is highly likely and that wildlife can serve as reservoirs as well as potential indicators of environmental contamination with AMR substances. The resistance to critically important antimicrobials (CIAs) found in this study as well as in other studies [24,25,34,35], which are considered as last resources for the treatment of complicated infections in human medicine [22], emphasizes the possible repercussions for both animal and human health, highlighting the need for stringent monitoring of antimicrobial use.

Due to their ability to travel great distances and cover diverse ecological niches, migratory birds have been recognized as powerful disseminators of AMR [5,23]. This study’s power to identify AMR in long-distance migratory birds was limited due to the small number of long-distance migratory birds tested in this study. Indeed, out of the two long-distance migrating birds sampled, we found no resistance. Although the bird species tested in this study were predominantly resident or short-distance migrants, these birds and other wildlife species still carry the potential to spread AMR throughout their range in diverse environments, including mountain regions and remote areas.

Birds associated with water are suggested to be likely to carry AMR bacteria [5,36], since water is known as important transmission medium for AMR [9,37] and AMR bacteria have been detected in various water samples [38]. Water-associated bird species examined in this study showed a higher AMR prevalence compared to non-water-associated bird species. However, due to the limited sample size of water-associated birds in this study, it is difficult to draw firm conclusions. We found no association between the location of the sampled wildlife regarding the percentage of water in the habitat and prevalence of AMR. This discrepancy raises questions, as higher values in AMR prevalence have been expected in wildlife sampled in areas with high water resources. A possible explanation for this discrepancy is that our habitat analysis covered only 1 km radius around the sampling sites. Most wildlife is very mobile and might have extended habitats well beyond this radius. Therefore, the sampling locations may just partially represent the actual habitat of these species. This highlights the complexity of studying the effects of specific habitat characteristics on AMR prevalence among wildlife. It underlines the need for more comprehensive studies to consider the full extent of wildlife mobility and the association with habitat.

Research exploring the association of feeding behavior with AMR prevalence in bacteria is limited. Although a few studies reported that feeding behavior may explain differences in AMR prevalence among different wildlife species, comprehensive investigations in this area remain (i) relatively rare [9,13,14], (ii) they typically enrolled only one wildlife species [11,14,25,27] or (iii) they involved species with similar feeding behavior such as wildlife with herbivorous or carnivorous diets [13,16]. This poses limitations on the attempt to directly compare wildlife species and their species-specific traits such as their feeding behavior. To address this limitation and to compare resistant *E. coli* bacteria in different wildlife species, our study sampled a broad range of wildlife species. The study revealed a significant association between feeding behavior and AMR prevalence. This is consistent with findings of a study, which indicated lower AMR prevalence among roe deer compared to wild boar [35]. In addition, another study showed that omnivorous wildlife species often move close to human settlements and farms, where they feed on animal waste or human garbage [25]. This feeding behavior increases their exposure to antimicrobials and bacteria of human origin. Accordingly, other studies suggest that herbivorous diets may lead to lower AMR prevalence due to microbiota dynamics and dietary adaptation to available resources [7,13]. Omnivores cover larger areas in search for food, often bringing them close to more densely populated areas, compared to herbivorous species that closely adapt to vegetation [7].

Available data frequently lack exact metrics describing the habitats of wildlife [9]. Furthermore, studies focused on a limited area for sample collection, thus providing localized insights rather than a comprehensive understanding of AMR prevalence across regions. For instance, one study exclusively focused on wildlife in one Italian national park, limiting the generalizability of their findings [16]. Similarly, studies investigated small wild mammals in England and Canada with only limited home ranges, making it challenging to infer general assumptions for broader areas [10,26]. Additionally, other studies examined wildlife from wildlife rescue centers, which limits comparability due to a lack of information about the wildlife’s origin [23,31,39]. Our study provides a transregional, comprehensive approach by including ecological features of the locations where wildlife species were sampled. By choosing this approach, we avoided limited interpretability of our data due to small sample sizes, having enrolled few or only very similar species or regional confinement. The findings revealed an association between a decreasing AMR prevalence and sample locations with higher percentage of forested area. Forests can serve as a retreat location for wildlife and the lower AMR prevalence may be largely attributed to less human interaction in this area. In the literature there is a strong suggestion by previous studies that the degree of human interaction is associated with the AMR prevalence in wildlife [12,33]. One example for such human activity are wastewater treatment plants [40]. The observed inverse association between the proportion of forested areas and the prevalence of AMR highlights a significant positive association of natural environments with lower presence of AMR bacteria in wildlife.

## 5. Conclusions

Our study presents a new large-scale quantification of AMR prevalence in wildlife across Switzerland that confirms the presence of AMR and MDR across a diverse array of wildlife species. We used this large and diverse data set to identify factors associated with AMR in wildlife species, particularly examining the impact of taxonomic differences, including feeding behaviors, and environmental characteristics. The observed AMR patterns in *E. coli* showed variability across different host species, as well as associations with environmental characteristics. These results underscore the association between the prevalence of AMR, environmental characteristics, and feeding behavior of wildlife. The detection of AMR in free-ranging wildlife underscores the extent of anthropogenic impact on natural ecosystems and underscores the necessity of an integrative ‘One Health’ approach to address this escalating public health concern. Understanding the connections of AMR is crucial given humans’ reliance on antimicrobials for curing bacterial diseases.

## Funding

This work was supported by the Novartis Foundation for medical-biological research [grant number 22A018]. B.Y. is supported with Swiss National Science Foundation (SNSF) Starting Grant (TMSGI3_211300) and SNSF Ambizione Grant (PZ00P3_185880). The funding agency did not interfere with studies design, survey, analysis and interpretation, or publication of the data.

## Competing interests

None declared.

## Acknowledgements

The authors thank all game wardens, hunters and researchers for their contribution to the project. We thank Christian Kühni of the Federal Office for Topography (swisstopo) and Beat Tschumi of the Federal Office for Agriculture (FOAG) for their valuable data provision.

## Ethical approval

Activities were approved by the competent authorities (permits P38 and BE61/2020/32505).

## Notes

### Competing Interest Statement

The authors have declared no competing interest.

